# Temperature and guanidine hydrochloride effects on the folding thermodynamics of WW domain and variants

**DOI:** 10.1101/2021.07.16.452632

**Authors:** Meng Qin, Natalia Denesyuk, Zhenxing Liu, Wei Wang, D. Thirumalai

## Abstract

We used simulations based on an all atom Go model to calculate the folding temperatures (*T*_*f*_s) and free energies (Δ*G*s) of two variants of the WW domain, which is a small all *β*-sheet protein. The results, *without adjusting any parameter*, are in good agreement with experiments, thus validating the simulations. We then used the Molecular Transfer Model to predict the changes in the Δ*G* and *T*_*f*_s as guanidine hydrochloride concentration is varied. The predictions can be readily tested in experiments.

## Introduction

In this paper, we study thermal denaturation and effects of guanidine hydrochloride (GdmCl) on the thermodynamics of folding of the well-characterized small all *β* WW domain^1–3^ and two variants using all atom Go (AAGO) model.^4–6^ Denaturation effects are accounted for using the successful Molecular Transfer Model (MTM).^7–10^ The rationale for using the Go model using lattice^11–14^ or off-lattice^15,16^ Go and AAGO models is that non-native interactions only play a sub-dominant role in determining the folding mechanisms of globular proteins.^17,18^ The computations based on the Go model provided support to theory, ^19–22^ which ushered in the notion that the gradient in the funneled energy landscape drives folding predominantly to the native state under folding conditions.

The studies cited above and subsequent developments (see perspectives by several authors^23–29^) have given tool sets, which have firmly established that small single domain proteins fold by parallel routes. Most of the computations, which have led to far reaching conclusions, have been done by triggering folding by changing the temperature. Experiments, on the other hand, typically investigate folding by altering the concentration of denaturants, such as urea or GdmCl at a fixed temperature. Therefore, practical ways of including denaturant effects are needed so that quantitative comparison between simulations and experiments can be made. We introduced the MTM to accomplish this goal.^7,30^ Currently, MTM is the only method that can be used to obtain fairly accurate results for denaturant-induced folding (see for example^8–10^), regardless of the size, sequence or topology of the folded state.

Our goal is to calculate the thermodynamic properties of the all *β* sheet protein using AAGO, and then predict how they change as function of GdmCl. The folded state of the wild-type Pin WW domain (PDB entry:1PIN), spanning Lys6 to Gly39, has a twisted triple-stranded antiparallel *β*-sheets (Figure 1). There are two loops. The first, loop1, has 6 residues (S_16_R_17_S_18_S_19_G_20_R_21_), and the shorter loop 2 has 4 residues. Another important member of the WW domain family, the Formin Binding Protein 28 (FBP28) WW domain (PDB entry: 1E0L), spanning residues from Ser6 to Pro33, has a structure that is similar to the 1PIN WW domain. Except for the difference in the 5 residue loop1 (T_13_A_14_D_15_G_16_K_17_), FBP28 is homologous to the 1PIN sequence. Not surprisingly, they have a similar folding behavior. If the 6-residual loop1 in 1PIN WW domain is mutated with the 5-residual FBP28 loop1 creates a variant (PDB entry:2F21) the thermodynamic stability is enhanced. The melting temperature increases from from 332K (1PIN) to 350.5K.^1^ Here, we focus on 1PIN and 2F21 domains.

**Figure 1:**
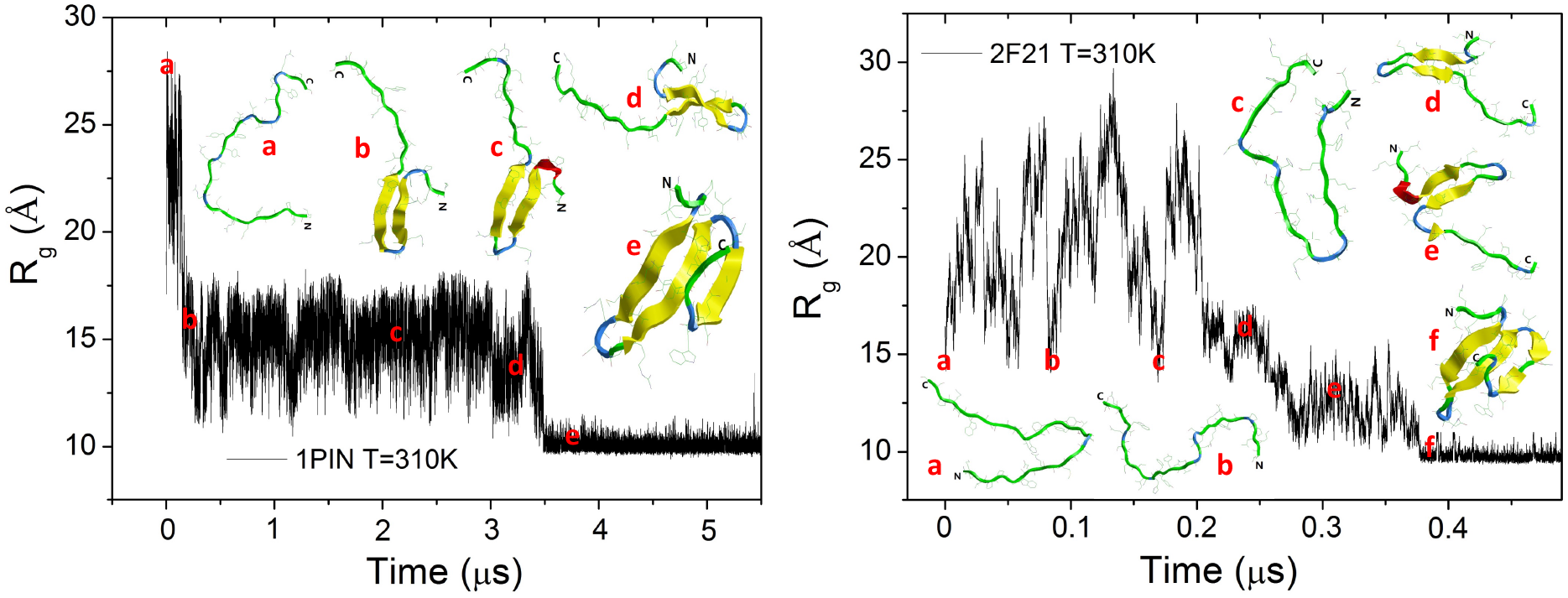
Folding trajectories at 310K for the 1PIN (2F21) domain on the left (right). The dynamics of folding is monitored using *R*_*g*_ as a function of time. Snapshots (a-e) and (a-f) are conformations that are sampled by 1PIN and 2F21, respectively. In both the trajectories a conformation with a *β*-hairpin ((c) in 1PIN and (e) in F21) form upon collapse of the proteins.

In all of our previous applications of the MTM we used a coarse-grained Self-Organized Polymer (SOP) model^31^ in which each residue in the protein was represented using two beads, one centered at the *α*-carbon and the other at the center of mass of the side chain. Here, we developed the AAGO model, following previous studies, ^4–6^ to model the WW domains. We used the MTM in concert with AAGO to investigate the effect of GdmCl on the folding of WW domain and two variants, after ensuring that the simulations in the absence of denaturants reproduce the experimental results quantitatively.^1,2^ We then predicted the thermodynamic properties for the two variants as a function of GdmCl. The current work shows that the tools used here are sufficient to anticipate denaturant effects on the thermodynamics of globular proteins.

## Model and Simulation Methods

### All-atom Go model (AAGO)

In the AAGO model all heavy (non-hydrogen) atoms are included, and bond lengths, bond angles, dihedral bonds are maintained as harmonic potentials. As is customary in Go models, there is an attractive potential between non-bonded atom pairs (heavy atoms *i* and *j*, where *i > j* + 2) that are in contact in the native state. All other non-local interactions are repulsive, which ensures that the atoms have a defined excluded volume.

Native non-bonded interactions separated by exactly three bonds (1-4 interaction) are reduced by a scale factor 0.5, in calculating the dihedral potentials. We used the standard functional form of the potential, given by,

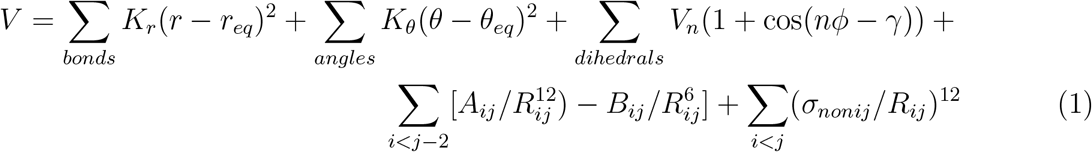

In the above equation, the atom types, and parameters for bond angle and dihedral interactions are taken from Table 14 given in the amber ff94 force field. ^32^ The equilibrium values of *r*_*eq*_, *θ*_*eq*_ values are taken from the X-ray structures of the folded proteins, and the *K*_*θ*_ value was taken from vibrational analysis of small molecules containing the specific atoms and standard normal mode analysis, as previously described.^32^ The force constants associated with the (*φ, ψ*) angles are chosen as specified in the Amber force field ff99SB (Table 1 in^33^). In the above equation, *V*_*n*_ is the amplitude of the dihedral angle potential, *n* is dihedral periodicity, and *γ* is the phase of the dihedral angle, *φ*. The Fourier series is approximated using a small number (*n* up to 3) of terms. The form of the improper torsions is the same as the dihedral angles.

In order to reduce the simulation time, the side chain of each amino acid is assumed to be rigid. This simplification means that the relative positions of these heavy atoms in one side chain are rigidly fixed during the simulations. The movement of these atoms in rigid body can be decomposed into the translational of the center of mass and the rotational movement of the rigid body. This approximation, which does not compromise accuracy, eliminates the need to calculate the bond stretch, bond angle, and dihedral angle potentials.

### Simulations

Extensive thermodynamic sampling of the protein WW domain is achieved using low friction Langevin dynamics^34^ simulations. The equations of motion for the position of a heavy atom, *r*_*i*_, are integrated using, 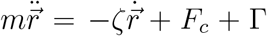 Here, *m* is the mass of the atom in each amino acid, *F*_*c*_ = −*∂V*/*∂r*_*i*_, and Γ(*t*) is a random force with a white noise spectrum satisfying the autocorrelation function, which in the discretized form reads *<* Γ(*t*)Γ(*t*+*h*) *>*= (2*ζk*_*B*_*T*/*h*)*δ*(0, *k*), where *δ*(0, *k*) is the Kronecker delta function, *h* is the integration step size, and *n* = 0, 1, 2, ….

We used the Verlet leapfrog algorithm to integrate the equations of motion. The velocity, *v*_*i*_, at time 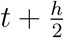 and the position, *r*_*i*_, at time *t* + *h* of an atom are given by,

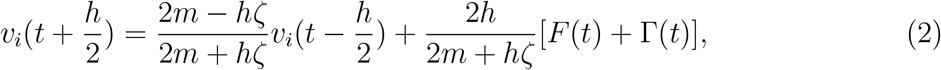

and 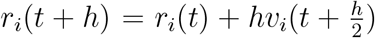 The value of the time step used to integrate above two equations is *h* = 0.02*τ*_*L*_, *τ*_*L*_ is the unit of time, and a low friction (a tenth of the viscosity in water) is used to obtain enhanced sampling and converged equilibrium thermodynamic data.

### Molecular Transfer Model (MTM)

In order to include denaturant effects, we used the MTM, which first involves computing the transfer free energy, which has been remarkably successful in studying the folding of a variety of proteins.^8,35^ The free energy of transferring a protein conformation, described by the set of coordinates *r*_*i*_s, from water ([*C*] = 0) to aqueous denaturant solution ([*C*] ≠ 0) is taken to be as *δG*(*r*_*i*_, [*C*]) = Σ_*k*_ *δg*(*k*, [*C*])*α*_*k*_(*r*_*i*_)/*α*_*Gly*−*k*−*Gly*_, where the sum is over backbone and side chain, *δg*(*k*, [*C*]) = *m*_*k*_[*C*] + *b*_*k*_ (*m*_*k*_ and *b*_*k*_, and *α*_*Gly*−*k*−*Gly*_ ^8^ are the transfer free energy of group *k, α*_*k*_(*r*_*i*_) is the SASA of *k*, and *α*_*Gly*−*k*−*Gly*_ is the SASA of the *k*^*th*^ group in the tripeptide Gly-k-Gly, respectively. The values of the slope *m*_*k*_[*C*] and the intercept *b*_*k*_, needed to calculate *δg*(*k*, [*C*]), are listed in Table S3 in a previous study.^8^ Thus, the effective free energy function for a protein at [*C*] ≠ 0 is *G*_*P*_ (*r*_*i*_, [*C*]) = *V*(*r*_*i*_) + *δG*(*r*_*i*_, [*C*]). We computed *α*_*Gly*−*k*−*Gly*_ by summing the contribution of all the SASA of the heavy atoms in the side chain and backbone.^36^ Note that MTM does not assume whether the protein folds by a two-state or multi-state mechanism.

It is important to realize that the MTM simulations could be performed by adding the *δg*(*k*, [*C*]) term to the energy function. Alternatively, if one is interested only in the thermodynamic properties then it suffices to sample the conformations in the absence of denaturants, and reweight them appropriately to account for the effects of denaturants.^8^ Elsewhere we have shown theoretically^9^ that the latter, more efficient procedure, is exact. In several previous studies, ^8,10,35,37^ we have shown that MTM and a coarse-grained model of a protein with beads accurately describes the thermodynamics and kinetics of folding of a number of proteins.

### Weighted Histogram Method

The heat capacity, *C*_*v*_ and the free-energy are calculated using the weighted histogram analysis method, which gives optimal estimate of the density of states of the system, allowing us to calculate the relevant thermodynamic quantities accurately. We calculated *C*_*v*_ and the free energy based on 25 independent long-time runs at different temperatures from 300K to 450K.

## Results

### Folding trajectories

Figure1 shows the folding trajectories of 1PIN and 2F21 domains. The Table of Contents graphics shows a folding trajectory for the FBP28 domain. Although the topology of the folded states are similar the pathways and the time needed to reach the native states vary greatly (see Figure 1). We plot the radius of gyration from an unfolded state for the two variants as a function of time, *t*. There is a precipitous decrease in *R*_*g*_ in ≈ 5*µs* as the protein folds from a fully denatured state. In the process of reaching the folded state, this WW domain samples a few intermediate states. The folding trajectory for the more stable 2F21 is different. In this case, only prior to reaching the folded state, the protein is compact. It is clear that in 2F21 folding, compaction occurs on similar time scales. In addition, the folding rate of 2F21 is about an order of magnitude greater than for the 1PIN. The ensemble of conformations sampled by 2F21 during the folding process is conformationally more heterogeneous than found for the less stable 1PIN.

### Heat capacity and the folding temperature (*T*_*f*_)

From the plots of *C*_*v*_ in Figure 2 as a function of *T*, we find there is a single peak for both 1PIN and 2F21 variants. The width of the transition is somewhat narrower for 2F21 than for 1PIN. The peaks in the *C*_*v*_ plots (in Figure 2), which we associate with the folding temperature, are in excellent agreement with experiments. This finding is particularly gratifying given that we did not *adjust* a single parameter to obtain agreement with experiments.

**Figure 2:**
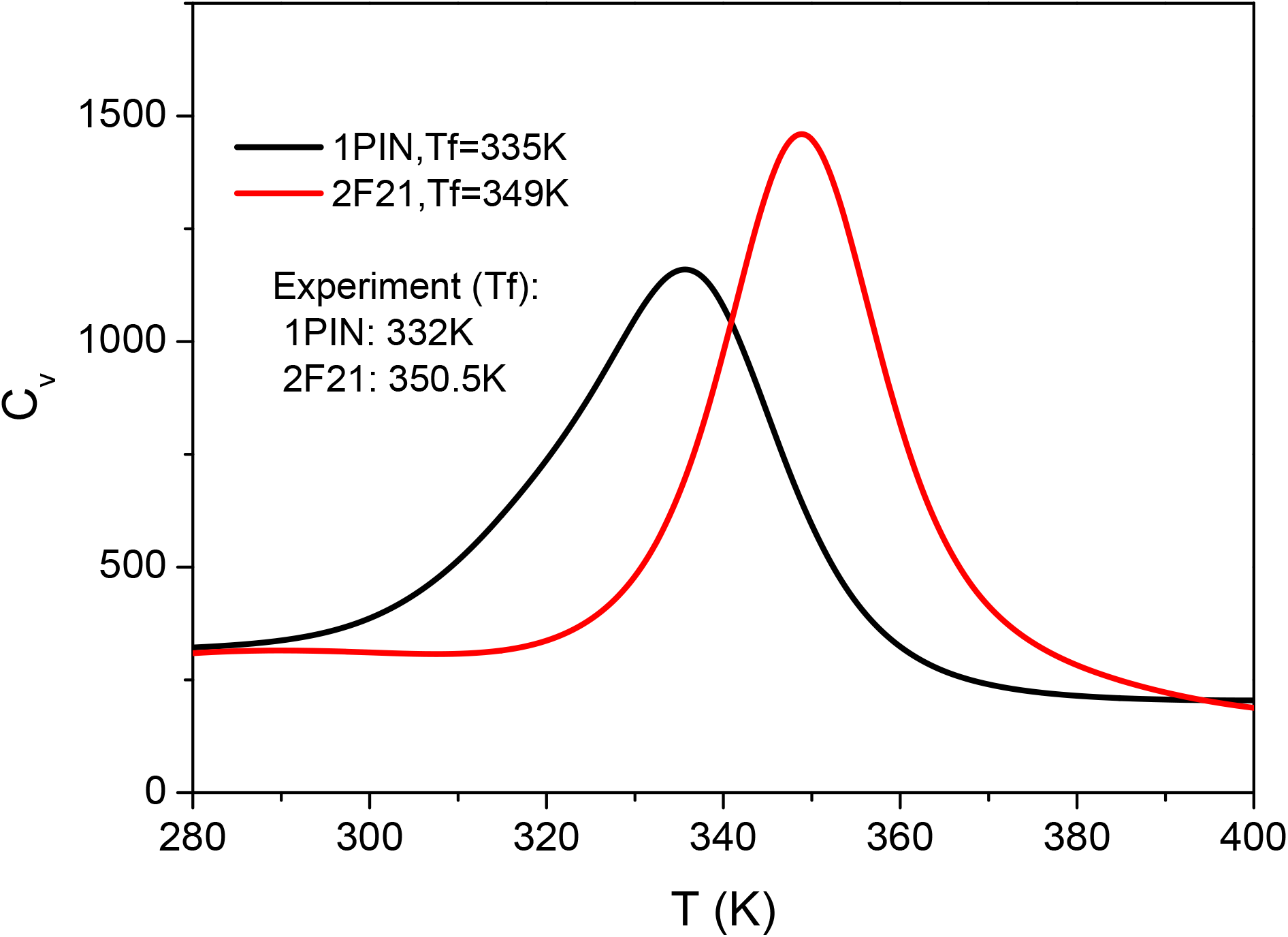
Heat capacities, *C*_*v*_, in units of in units of *cal/mol · K* for 1PIN (black) and 2F21 (red) as a function of temperature. The folding temperatures correspond to the peaks in *C*_*v*_. The experimental values are also shown.

### Temperature-dependent Energies

Heat capacity *C*_*v*_ (in Figure 2) as a function of *T* shows that 1PIN and 2F21 domains fold in a two state manner. Therefore, we could compute the difference between the folded and unfolded states using, 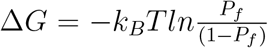 where *P*_*f*_ is the probability of being in the folded state. A given conformation if the structural overlap (see Eq. S16 in a previous study^8^) with the folded state is small. Comparison of the calculated and experimental values for Δ*G* as a function of *T* for both the variants shows very good agreement (Figure 3). Note that Δ*G* = *G*_*u*_ − *G*_*f*_ where *G*_*u*_ and *G*_*f*_ are the free energies of the unfolded and folded states, respectively.

**Figure 3:**
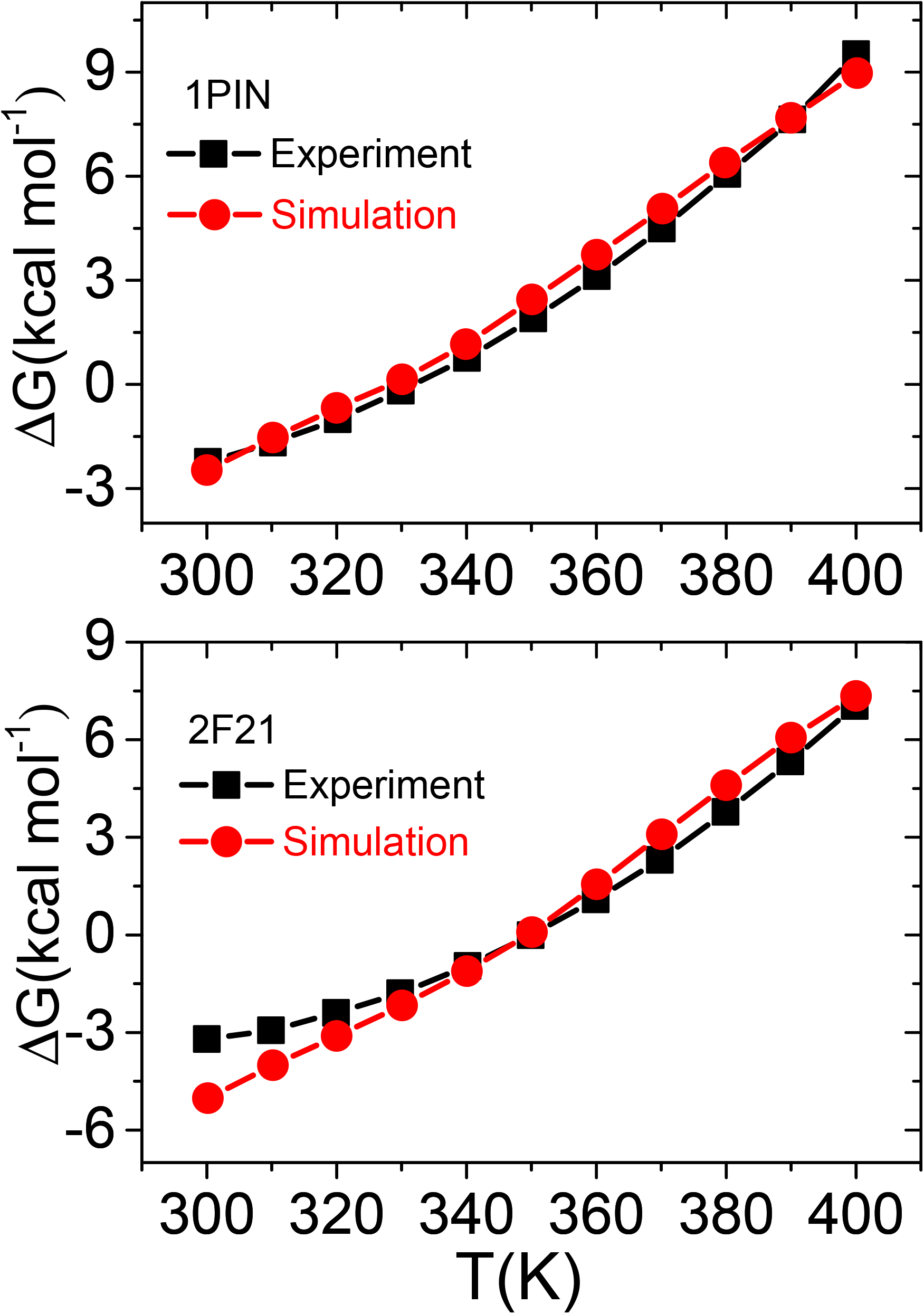
Free energy difference between the folded and unfolded states as a function of temperature. Top and bottom panels are for 1PIN and 2F21, respectively. Black (red) lines are from simulations (experiments).

### GdmCl effects on folding thermodynamics

The results in Figures 2 and 3 show that the all atom Go model reproduces the folding thermodynamics as a function of temperature, thus validating our model. We used the ensemble of conformations generated at various temperatures to calculate the GdmCl-dependent folding temperature and free energies using the MTM model. Plots of *C*_*v*_ at various values of the GdmCl concentration are displayed in Figure 4. Two observations can be made: (1) The dependence of the folding temperature on GdmCl changes linearly as the concentration of the denaturant increases. (2) At all values of the denaturant concentrations *T*_*f*_ for 2F21 is greater than for 1PIN, which is refection of the differences in the stabilities. A similar conclusion may be reached from the plot of Δ*G* as a function of GdmCl concentration at the indicated temperatures (Figure 5).

**Figure 4:**
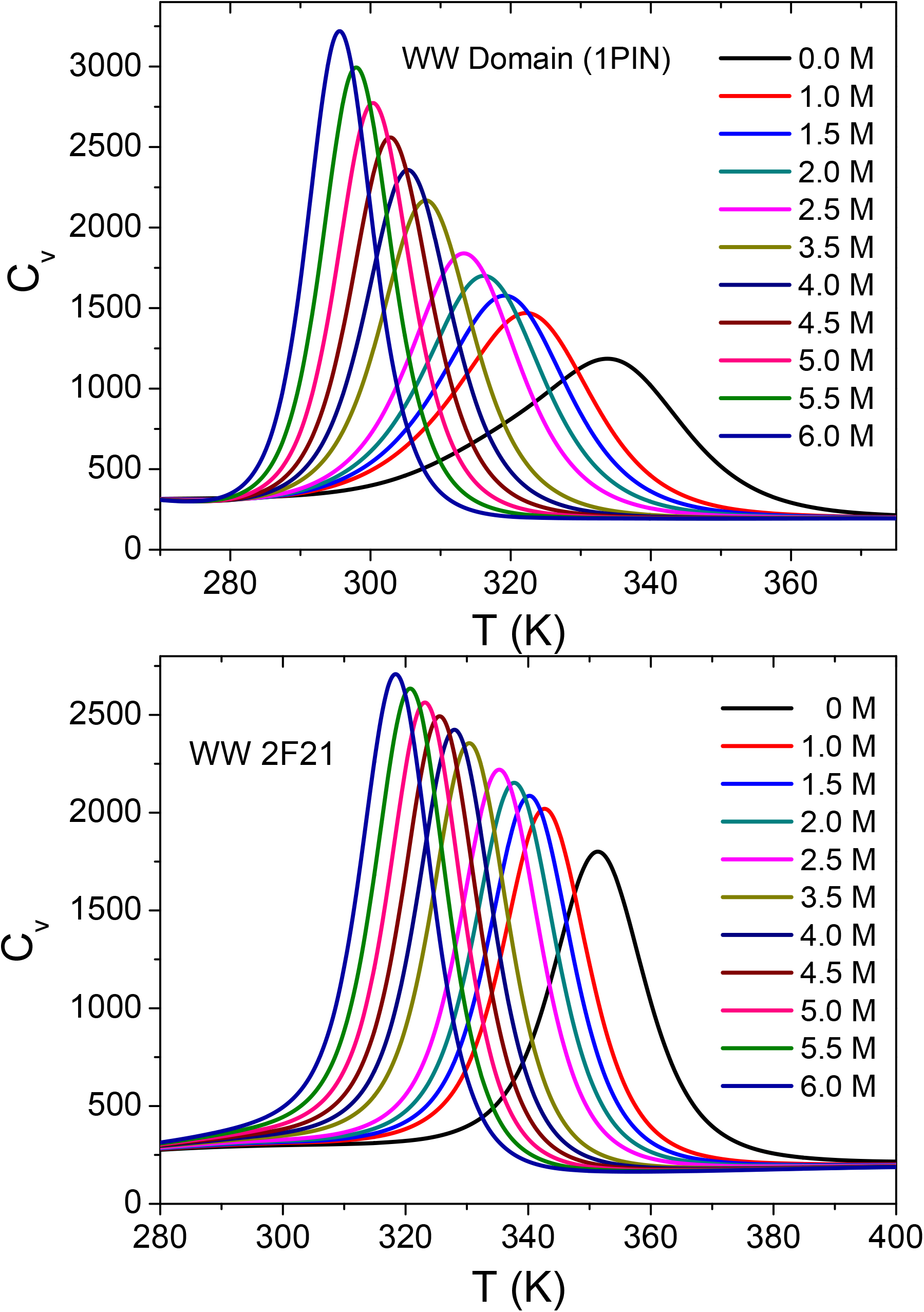
The dependence of *C*_*v*_, in units of *cal/mol · K*, on temperature at various values of the GdmCl concentration. Top and bottom panels are for 1PIN and 2F21, respectively. The two-state transition at all values of denaturant concentration is reflected in a single peak in the heat capacity.

**Figure 5:**
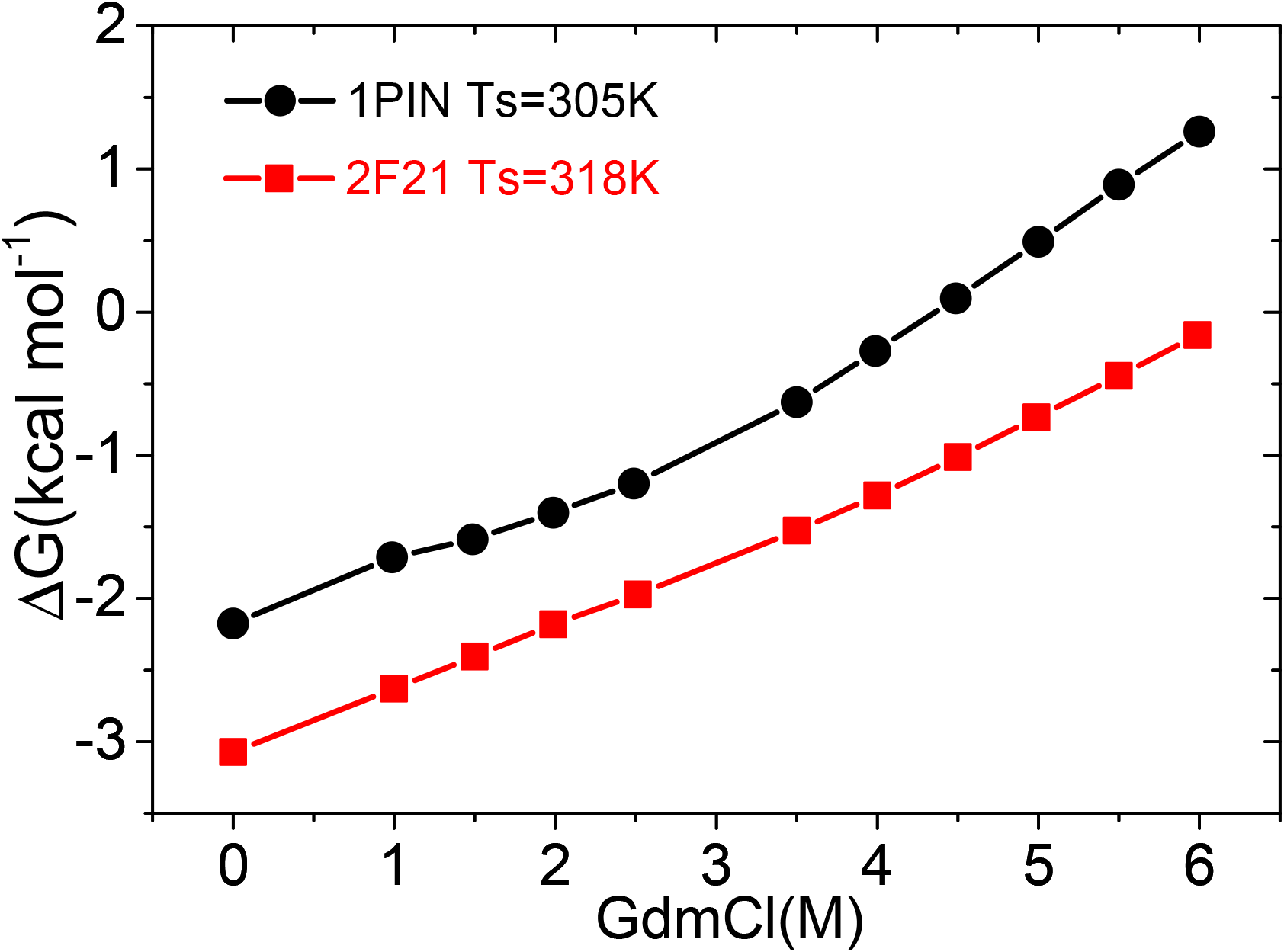
Predicted free energy difference between the folded and unfolded states as a function of GdmCl for 1PIN(black) and 2F21(red), respectively. The simulation temperatures are shown.

## Discussion and Conclusion

We have reported the folding thermodynamics of two variants of a small all *β*-sheet protein. The good agreement, without adjusting any parameter to fit measurements, between simulations and experiments for thermal folding shows that AAGO is a reasonable starting point for investigating the folding of globular proteins. The results for GdmCl effects, shown in Figures 4 and 5, are genuine predictions, which could be readily tested.

### Additional comparison to Experiments

In a previous experimental study,^38^ the thermodynamic properties of five sequences, all with the topology resembling the WW domain, were investigated using different probes. Although the sequences are different from the ones we simulated here, there are a few results that are consistent with our findings. (1) The folding temperature (referred to as the melting temperature in the experiments) correlated with stability, which is clearly what is expected. (2) More interestingly, the folding temperature varies linearly with the denaturant concentration. In the experiments urea was used as the denaturant but here we used GdmCl in which electrostatic interactions play a role.

The linear dependence of the stability changes calculated using simulations (Figure 5) is in qualitative agreement with experimental data^39^ on Fip35 WW domain. Figure 2 in Wirth and Gruebele^39^ shows that the both *lnk*_*u*_ and *lnk*_*f*_ change linearly as a function of GdmCl concentration. This implies that *DeltaG* must increase linearly as GdmCl is increased, which is consistent with our simulations. It is also gratifying that the time scales of folding in the experiments (on the order of tens of *µs*) also is supported in our simulations.

### Limitations

The calculation of the folding temperatures for the WW sequences, which are in excellent agreement with experiments, using the Go model with standard Amber force field is encouraging. In addition, we have determined the dependence of *T*_*f*_ on the concentration of GdmCl, which is in qualitative agreement with changes in the melting temperature as a function of urea for five unrelated WW sequences. However, there are points of disagreement between simulations and experiments.^38^ The most glaring is the that the heat capacity changes calculations in the simulations do not agree with experiments. More worrisome is the observation that the sign of the calculated heat capacity (Δ*C*_*v*_) is the opposite of what is found in experiments even though the calculated and measured numerical values of Δ*C*_*v*_ are small for WW domains. ^38^ We believe the main reason is that hydration effects, especially in the unfolded states, are not modeled correctly in the coarse-grained models. Although it is unclear how one could solve this issue without sacrificing the simplicity of the coarse-grained models, this has to be fixed if one were to calculate Δ*C*_*v*_ accurately.

### Folding cooperativity

As stated earlier, the calculated dependence of the folding temperature on denaturant agrees qualitatively with measurements.^38^ It is interesting to examine the sequence dependendent variations in cooperativity in WW domains. Assessing the extent of cooperativity is not straightforward because one needs to use a precise mathematical definition. To make the situation more complicated, one finds that even *T*_*f*_ depends on the probe. In the context of the WW domains, this is illustrated vividly in Figure 2 in the experimental study. ^38^ This figure also illustrates visually that the sharpness (or extent of cooperativity) could vary as the probes (CD or FT-IR) are changed. Thus, a quantitative measure for calculating or measuring the extent of cooperativity, which could be used in experiments and simulations, is needed. Elsewhere we introduced one such measure, Ω_*c*_.^40^ We showed that for two state folders Ω_*c*_ ∝ (Δ*G*)^2^ (see Eq. 15 in^40^). Using this measure one would predict that folding cooperativity ought to be higher in 2F21 compared to 1PIN. This is indeed the case (see Figure 2). Because Ω_*c*_ ∝ (Δ*G*)^2^ it follows that *T*_*f*_ of 1PIN should be lower than 2F21. The results in Figure 2 confirm this expectation.

The values of Ω_*c*_ can be readily calculated using the data in Figure 2 in the experiment. ^38^ Alas, the sharpness of the transition depends on the probe used to infer the dependence of the probability of being folded as a function of temperature. For instance, by visual inspection we would expect that Ω_*c*_ should be the smallest for the orange curve for FBP11WW1 (Figure 2 in^38^) and be comparable for the black and green curves, which are obtained using different probes. This illustrates the essential difficulty in assessing the extent of cooperativity using experimental data. Nevertheless, it is clear from Table 2 that the sharpness of the transition tracks stability, just as in the simulations. It could be useful to make a quantitative comparison by calculating Ω_*c*_ using experimental data. ^41^

### Final remarks

(1) The all atom Go model by definition ignores non-native interactions. This is a limitation of the model even though we have reproduced the experimental results accurately (Figures 2 and 3). It is necessary to include non-native interactions for some test cases in order to assess the errors, if any, in the calculation of folding thermodynamics. (2) Hydration effects are completely neglected and could also be important in describing the equilibrium aspects of folding. One way of accounting for hydration effects is to simulate folding of all atom Go model in explicit solvent. If this protocol is successful then the role of water can be directly addressed. The inability of this model or previous lower resolution coarse-grained models^8^ to correctly reproduce the heat capacity differences between the folded and unfolded states is directly attributable to the absence of proper treatment of hydration effects, which affects the unfolded state more than the folded state. (3) Given the success of the MTM, we believe it will be interesting to use the conformations generated using atomic detailed simulations for WW domain^3^ and use them in conjunction with MTM to obtain the effects of denaturants (GdmCl or urea) on the folding thermodynamics. Perhaps the most important conclusion of our study is that the all atom GO model using the standard form for the energy function and MTM may be used to obtain the thermodynamic properties of proteins in the presence of denaturants. We conjecture that if the model could be simulated in water, it is likely that the asymmetry observed in experiments in the temperature dependence of heat capacity may be recovered. If this proves to be the case then our computational framework could prove to be be sufficiently accurate in calculating the effects of denaturants on the folding thermodynamics of proteins.

## Acknowledgement

We are also grateful to George Makhadatze, Martin Gruebele, and Angel Garcia for useful discussions. Most of the work was done while MQ and DT were at the University of Maryland. This work was supported in part by a grant from National Science Foundation (CHE 19-00093) and the Collie-Welch Chair (F-0019) administered through the Welch Foundation. ZL acknowledges financial support from the National Natural Science Foundation of China (11104015,11675017,11735005).

